# Algorithms for Search for Common Ancestors

**DOI:** 10.1101/238188

**Authors:** Rita Gitik

## Abstract

Generation theory was developed as a tool for studying self-reproducing systems. In this paper we show that this theory can be applied to a search for common ancestors of living organisms.

We give two algorithms with flowcharts in pseudocode for finding common ancestors of a set of microbes and describe the connections with the genography problem.

## Introduction

This paper describes certain aspects of Genography, which proposes on the basis of fossil and genetic evidence to make specific statements on the origin of groups and species. Examples of such statements are: *all modern members of group X living in location L descend from N individuals that lived Y years ago in location M*, or *species X and species Y have a common ancestor from which they branched Z years ago*. An instance of the first statement is implied in [13], which claims that all European humans are descendants of seven women who lived in Africa 45,000 years ago. An instance of the second statement is implied in [14], which claims that gorillas and chimpanzees have a common ancestor, from which they branched around 5,000,000 years ago.

Genography is motivated by a deep-seated human search for the origin of species. In the case of humans, IBM and the National Geographic Magazine are running the Genographic project, about which information is available at the National Geographic website [16]. The study published by the Genography Consortium in [1] contains extensive bibliography supporting the project.

However, in spite of the numerous appearances in popular science publications, Genography has not received rigorous scientific scrutiny. Even though there is a number of papers criticizing the genography project on moral and ethical grounds (see, for example, [15]), the author had not been able to find scientific analysis of the project in print. The purpose of this paper is to fill this gap.

We exhibit a careful mathematical analysis of the plausibility of the basic claims made by Genography and show that the problem is mathematically ill-posed. This analysis is novel and constitutes the original contribution of this paper. The analysis is especially interesting in view of a current discovery, reported in [5], of another new hominin species, namely *australopithecus deyiremeda*, which lived about 3.5 millions years ago.

In the nineteenth century, mathematician Jacques Hadamard defined a well-posed problem as a mathematical model of physical phenomena with the following properties:

1. A solution exists.
2. The solution is unique.
3. The solution depends continuously on data in some reasonable topology.

Problems that are not well-posed in the above sense are said to be ill-posed. Well-posedness is an important technical concept because data are often corrupted by random measurement errors. Hence, when a problem is ill-posed, such measurement errors, even though they may be small, can cause large changes in the solution, making it meaningless. On the other hand, when a problem is well-posed, its solution can be reliably computed using a stable numerical algorithm, in spite of small measurement errors in the data. A common method for dealing with ill-posed problems is to reformulate them with additional assumptions in a process known as regularization, to make them well-posed.

The technical approach of this paper uses Generation Theory, which was introduced in [7] to analyze self-reproducing systems. Within the framework of Generation Theory the following original results are obtained about Genography:

1. Genography is ill-posed.
2. It is possible to regularize the original Genography problem so that it becomes well-posed.
3. We exhibit two algorithms for the solution.

The remainder of this paper is as follows: we present an overview of Generation Theory and introduce a problem, called *Microbes in a jar*, which shows that Genography is ill-posed. Then we formulate a regularized *Microbes in a jar* problem, and give two algorithms with flowcharts in pseudocode for solving it.

## Background in Self-Reproducing Systems

The paper [7] is devoted to a study of self-reproducing systems. The idea goes back to von Neumann, who discussed cellular automata capable of building other cellular automata in [9]. The subject proved to be a fruitful ground for research. For a survey of existing literature and current developments see [3], [8], [10], and [12]. In the framework of generation theory, the entities that can potentially reproduce are called *machines*, regardless of their physical nature (e.g. robots, microbes, or lines of computer code). Reproduction is achieved by the action of a machine on available resources, producing an outcome that may or may not be a machine itself.

A generation system is defined as a quadruple (*U, M, R, G*), where *U* is a universal set that contains machines, resources, and outcomes of attempts at self-reproduction. *M* ⊆ *U* is a set of machines, *R* ⊆ *U* is a set of resources that can be used for self-reproduction, and *G*: *M* × *R* → *U* is a generation function that maps a machine and a resource into an outcome.

The generation sets are defined as follows: *M*_0_ = *M* is the set of all machines and *M*_*i*+1_ is the set of all machines that are capable of producing a machine in *M_i_*.

It was shown in [7] that the generation sets are nested, i.e. *M*_0_ ⊇ *M*_1_ ⊇ *M*_2_ ⊇ ⋯, which leads to the definition: *M*_∞_ = ∩*M_i_*. All self-reproducing machines belong to *M*_∞_.

In this paper we show that generation theory can be applied to study reproduction of microbes and describe the connections with the genography problem.

## Levenstein metric

For our purposes a living organism is represented by its DNA. We assume that a DNA is a finite sequence of symbols chosen from the alphabet of 4 letters: *A, C, G*, and *T*.

Levenstein metric *ρ*, which is a generalization of the Hamming metric, on a set of DNA is defined as follows.

1. For any two sequences of equal lengths *S* = (*s*_1_, ⋯, *s_n_*) and *R* = (*r*_1_, ⋯, *r_n_*) define *ρ*(*S, R*) = 0 if *s_i_* = *r_i_* for all 1 ≤ *i* ≤ *n*. Define *ρ*(*S, R*) = 1 if the sequences differ in exactly one coordinate. Define *ρ*(*S, R*) = *m* if the sequences *S* and *R* differ in exactly m coordinates.
2. If *S* can be obtained from *R* by either deleting or inserting one coordinate, define *ρ*(*S, R*) = 1.
3. If the sequence *S* can be obtained from *R* by changing *k* coordinates and deleting or inserting *m* coordinates, define *ρ*(*S,R*) = min(*k* + *m*), where the minimum is taking over all possible choices of *k* and *m*.

**Example 1**. *Let S* = *ACCGCA and R* = *CTGA. By observation, the shortest way to obtain R from S is to delete the first and the fifth letters in S and to change the third letter from C to T. Hence ρ*(*S, R*) = 3.

It is easy to see that for any sequences *S* and *R*, *ρ*(*S, R*) = 0 if and only if *S* = *R*. Moreover, as *ρ*(*S, R*) = *ρ*(*R, S*), it follows that *ρ* is symmetric.

Note that *ρ* satisfies the triangle inequality. Indeed, if *S, R*, and *T* are sequences with *ρ*(*S, R*) = *m* and *ρ*(*R, T*) = *n*, we can transform *S* into *R* in *m* steps, and *R* into *T* in *n* steps, where a step consists of either deleting or inserting a coordinate, or of changing a coordinate. Hence we can transform *S* into *T* in *m* + *n* steps, therefore *ρ*(*S, T*) ≤ *m* + *n*, proving the triangle inequality. So *ρ* is, indeed, a metric on the set of DNA sequences.

Note that if *S* and *R* have equal lengths, then *ρ*(*S, R*) is the Hamming distance between *S* and *R*.

The motivation for the use of the Levenstein metric comes from the fact the two most common types of mutations in DNA are deletion or insertion of a letter in the sequence, or substitution of a different letter for a given one. The first type of mutation is called a frame-shift mutation and the second type of mutation is called a point mutation. The letters in the DNA sequences are called the nucleotides. For more information on mutations see [4] and [6].

To simplify the exposition we would like to define an induced semi-metric on the set of living organisms by *ρ*(*O*_1_, *O*_2_) = *ρ*(*DNA*(*O*_1_), *DNA*(*O*_2_)), where *O*_1_ and *O*_2_ are any organisms. This is a semi-metric because distinct organisms can, a priori, have identical DNA.

## Microbes in a jar

Consider a sealed jar containing liquid, whose chemical composition and temperature do not depend on time. This liquid is the set of resources *R*, so *R* is a singleton. Let *M* be the set of DNA of all microbes, i.e. one-celled organisms, which ever lived in this jar. *M* is the set of all machines in the problem. *M* has a decomposition *M* = *M^a^* ∪ *M^d^*, into subsets of alive and dead microbes. We define a generation function *G*: *M^a^* × *R* → (*M^d^* × *M* × *M*) by *G*(*m^a^, r*) = (*m^d^, m*_1_, *m*_2_). So the generation process takes a living microbe *m^a^* and produces, by division, two new microbes *m*_1_ and *m*_2_. The original microbe *m^a^* dies in the process, so the output of the generation function contains *m^d^*, to remind us that the original microbe does nor survive the reproduction. We require the following property.

## The Small Mutations Property

Assume that there exists a constant *ϵ* = *ϵ*(*M*) > 0, which does not depend on a particular *m* in *M*, such that if *G*(*m,r*) = (*m^d^*, *m*_1_, *m*_2_) then *ρ*(*m*, *m*_1_) < *ϵ* and *ρ*(*m*, *m*_2_) < *ϵ*. This requirement reflects an accepted empirical fact that in the absence of radiation mutations are not large. For example, such mutations cannot produce a two-celled organism, so an offspring of a microbe is a microbe. Note that two different microbes might have identical DNA, and as a consequence, we cannot identify microbes with their DNA, so any metric on the DNA will be only a semi-metric on the set of microbes. It implies that identical twins and genetic loops are theoretically possible. (A genetic loop will appear if a sequence of generations starting from some machine *m* will produce *m*.)

## Problem formulation and a connection with Genography

We want to know how many microbes were in the set *M*\* that generated *M*, what was their genetic makeup, and how many generations passed between *M*\* and *M*. We are given the set *M^a^* of all microbes currently alive and a subset *M^f^* of *M^d^*, which is a set of fossils. *M^f^* is usually very small relative to *M^d^*. In addition, for another small subset *M*′ of *M* we are given the set *G*^−1^(*M*′, *R*), so *M*′ is the set of microbes for which we know the parent. This is exactly the description of the genography problem. We know humans that are currently alive in a certain location, we know some of their ancestors, we know some of the family trees, and we want to obtain data about the first humans in this location.

### Remarks

1. Only alive microbes can reproduce, however their offspring might die at birth.
2. In this simple model we allow a microbe to die either at birth or after reproduction. No microbe survives reproduction.
3. *M*\* can be characterized as follows: *m* ∈ *M*\* ⇔ {*m* has no parent in the jar}.

Note that the length of the DNA of a virus is about 4 · 10^4^ nucleotides, for the microbe E. coli it is about 4.7· 10^6^, and for humans it is about 3· 10^9^. Under typical laboratory conditions E. coli reproduces every 20 min.

For E. coli the number of errors in DNA per replication under normal laboratory conditions before the error correction code starts working is about 1. The error correction code is very efficient and under normal laboratory conditions it results in one mistake per about 1000 replications, so a single mutation appears every 2 days.

## An Approach to the Solution

This problem is a typical example of reverse modeling, when we are given conditions in the present, and need to reconstruct the conditions in the past. It is well-known that the usual statistical methods are reasonably reliable only for short time intervals, so the first task is to estimate the time interval involved. Assume that the original microbes belong to a single species, so they reproduce at the same time interval *T*. Recall that the diameter of a set *X* is defined as *diam*(*X*) = sup*ρ*{(*x_i_,x_j_*)|*x_i_,x_j_* ∈ *X*}.

We assume that the small mutation property holds.

Let *Diam*(*M*\*) = *μ* ≥ 0, and let *ϵ* be as in the definition of the small mutation property. Then if *m*_1_ and *m*_2_ are microbes in the jar, and *n*_1_ and *n*_2_ are their respective parents, the triangle inequality implies that *ρ*(*m*_1_, *m*_2_) ≤ *ρ*(*m*_1_, *n*_1_) + *ρ*(*n*_1_, *n*_2_) + *ρ*(*m*_2_, *n*_2_). The small mutation property states that *ρ*(*m*_1_, *n*_1_) < *ϵ* and *ρ*(*m*_2_, *n*_2_) < *ϵ*, hence with each generation the diameter of the set of microbes in the jar increases by at most 2e. So the diameter of the *K*-th generation is bounded by *μ* + *K* · 2*ϵ*.

Determine all the pairwise distances between the elements of *M^a^* and, consecutively, the diameter of *M^a^*.

If *diam*(*M^a^*) = *μ* + *J* · 2*ϵ*, then the lowest bound on the number of generations in the jar is *J*. Hence the original population lived at least *J* · *T*[time units] ago, where *T* is the time interval between microbe divisions. (For example, *T* = 20 min for amebae.)

This estimate is very crude and it is easy to build models for which it is arbitrarily poor.

**Example 2**. *If each replication produces one dead microbe and one alive microbe with the exact DNA of the parent, then diam*(*M^a^*) = *diam*(*M*\*) = *μ, and the above estimate implies that J* = 0, *however, the number of generations may be arbitrarily large*.

So Example 2 illustrates that genography is ill-posed.

**Example 3**. *Assume that the replication is exact, and M^a^ contains exactly* 8 *microbes with identical DNA. We cannot decide whether M*\* *consisted of* 4 *microbes and J* =1, *or M*\* *contained one microbe and J* = 3.

Example 3 shows once more that genography is ill-posed and that we need to make a guess either about *M*\* or about *J*, or about both.

Assume that we have established the values of *J* and of *diam*(*M*\*).

In order to describe the set *M*\*, we assume that it is located in the center of the set *M^a^* and that *M^a^* is approximately a metric ball. Statistically speaking, we assume that all mutations have equal probability and that the lethal mutations are random.

A naive way to find the center of *M^a^* is to consider a pair of elements *m*_1_ and *m*_2_ such that the distance between them equals to the diameter of *M*^a^, and find an element or elements in *M^a^* whose distance to each of *m*_1_, *m*_2_ is closest to the half of the diameter of *M^a^*. This procedure would usually produce several answers. The next example shows that, unfortunately, the possible centers obtained in this way might be as far apart as the original points.

**Example 4**. *Consider the sequences* (0000) *and* (1111). *As ρ*((0000), (1111)) = 4, *ρ*((0011), (0000)) = 2, *and ρ*((0011), (1111)) = 2, *the sequence* (0011) *is half-way between* (0000) *and* (1111). *However, ρ*((1100), (1111)) = 2, *and ρ*((1100), (0000)) = 2, *so the sequence* (1100) *is also half-way between* (0000) *and* (1111).

*But ρ*((1100), (0011)) = 4.

Assume that we have found possible centers of the set *M*. Run *J* iterations of the generating function *G* on a metric ball of *diam*(*M*\*) around each candidate center, then check which output fits the set *M^a^* best. Alternatively, consider the frequencies of the DNA in *M^a^* and assume that the most frequent ones define the center. However, it is possible that each DNA in *M^a^* is carried by only one microbe. Then we might want to check DNA frequencies of small balls in *M^a^*, but at the end we will need to make a guess about M^*^ and to run forward simulations.

**Remarks**.

1. The sets *M^f^* and *M*′ are used to check the model. They are useful in estimating *J*. It will be shown in the next section that if these sets are maximal, then the problem has an exact solution.
2. As the accuracy of the statistical methods involved declines when *J* grows, many genography-type results are not widely accepted by the scientific community.
3. The assumption that all mutations have the same probability to be lethal is unrealistic. It seems that the lethal mutations should be localized in certain directions.

### Regularization of the Problem

In order to regularize the problem we need to introduce a very strong assumption.

## Condition UI (Uniqueness of Individual)

We assume that *ρ*, defined above, is a metric on the set of living organisms, and not just a semi-metric, i.e. *ρ*(*O*_1_, *O*_2_) =0 ⇔ *O*_1_ = *O*_2_, so the function from the set of organisms to the set of DNA is injective. Alternatively, we can define an individual as an equivalence class, but such approach is rather non-intuitive.

We consider the same setting as before, namely, there exists a jar in the laboratory which contains liquid, and the chemical composition and temperature of the liquid do not depend on time. This liquid is the set of resources *R*. Let *M* be the set of DNA of all microbes which ever lived in this jar. However, we demand that the system satisfies condition UI. Consecutively, to simplify the exposition, we identify a DNA sequence with its carrier.

We consider the same set of machines and the same generating function, *G*: *M^a^* × *R* → (*M^d^* × *M* × *M*) given by *G*(*M^a^, r*) = (*m^d^, m*_1_, *m*_2_). Note that condition UI implies that *m^d^* ≠ *m*_1_ ≠ *m*_2_.

We want to identify the set *M*\* of microbes which started *M*, i.e. the first microbes which appeared in the jar.

The following approach to the solution of the problem was developed by the author in [2].

Assume we know all the microbes which ever lived in the jar and their offspring. To be precise, we are given the set *M* = {*m*_1_, ⋯, *m_N_*}, as an ordered list. We also are given the set *r*(*M*) = *G*(*M, R*) as the ordered list of *L* pairs {(*m_i_*, *n_i_*)} G *M* × *M*, ordered by the first coordinate, where *m_i_* is a parent of *n_i_*. As each microbe in the jar has either 2 offspring or no offspring at all (the latter happens for microbes in *M^a^*), any *m_i_* ∈ *M* can appear as the first coordinate of a pair on the list *r*(*M*) exactly twice. As each microbe in the jar has at most one parent in the jar (the microbes without a parent in the jar are exactly the set *M*\*), any *n_i_* ∈ *M* can appear as the second coordinate of the list at most once.

We solve the problem by constructing the following graph Γ: the set of vertices of Γ is *M* and there is a directed edge from *m*_1_ to *m*_2_ if and only if *m*_1_ is a parent of *m*_2_ . Recall that the degree of a vertex in a graph is the number of edges attached to it. Also recall that a graph is a tree if it does not contain loops.

Assume Γ is constructed. As condition UI implies that there are no genetic loops, Γ is a forest, i.e. a collection of disjoint trees. Note that if m is alive, it has not reproduced yet, so for any live microbe deg(*M^a^*) = 1. In other words, the only edge attached to any living microbe in Γ is the edge connecting it to its parent. The same is true if m was born dead, thus could not reproduce. Also note that deg(*m*) = 2 ⇔ *m* ∈ *M*\*, because only in this case *m* has 2 offspring, but no parent in the jar. In all other cases deg(*m*) = 3.

We offer two algorithms for the construction of Γ. The algorithms will output the set *M*\* as an ordered list.

### Solution 1: Going forward in time

This solution does not use the small mutation property and is very general. Choose any *m*_1_ ∈ *M*, check who are its offspring, if any, and connect *m*_1_ to each of its offspring by a directed edge. Repeat this procedure with the offspring of *m*_1_. At each step the number of offspring may grow exponentially, but still there are only finitely many of them. After a finite number of iterations we will stop, constructing a graph Γ_1_ . If the set of vertices of Γ_1_ is *M*, then Γ_1_ = Γ. In this case Γ is a tree, and *M*\* = *m*_1_ . Otherwise, choose *m*_2_ which is not a vertex of Γ_1_ and repeat the process. After a finite number of steps we construct Γ, and we know that deg(*m*) = 2 ⇔ *m* ∈ *M*\*.

Below is a flowchart of this algorithm in pseudocode, where *L* is the length of the list *r*(*M*), and (*m_i_*, *n_i_*) are elements of *r*(*M*).

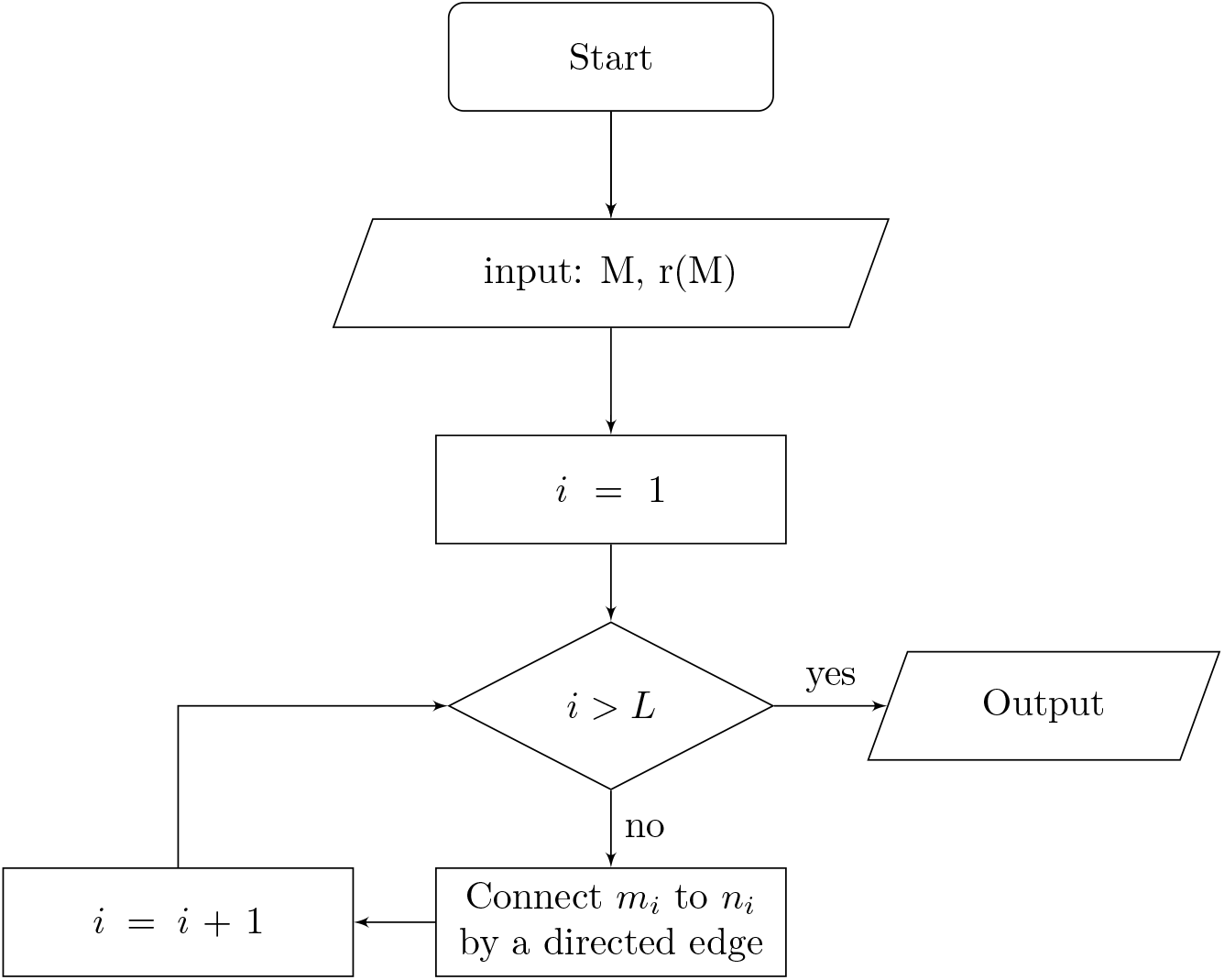

### Solution 2: Going backward in time

This solution uses the small mutation property. We assume that we can generate balls of radius *ϵ* in *M*, meaning that for any *m_i_* ∈ *M* we are given an ordered list *B_i_* of all *b_ij_* in an *ϵ*-ball around *m_i_*. Clearly the length of each *B_i_* is bounded by the length of *M*. Choose any *m*_1_ ∈ *M* and check who is its parent. The small mutation property implies that the parent belongs to the list *B*_1_, so to find the parent we need to check the offspring of all microbes in *B*_1_. Connect the parent to *m*_1_ and repeat the step with the parent of *m*_1_. The process terminates after finitely many steps, when we arrive at a microbe without a parent. Such microbe is an element of *M*\*. Call the graph constructed in the process Γ_1_. If Γ_1_ = Γ, then the set *M*\* is a singleton. Otherwise, choose *m*_2_ which is not a vertex of Γ_1_ and repeat the process. After a finite number of steps we construct Γ, and we know that deg(*m*) = 2 ⇔ *m* ∈ *M*\*.

Below is a flowchart of this algorithm in pseudocode, where *L* is the length of *r*(*M*), and *b_i_j__* ∈ *B_i_*.

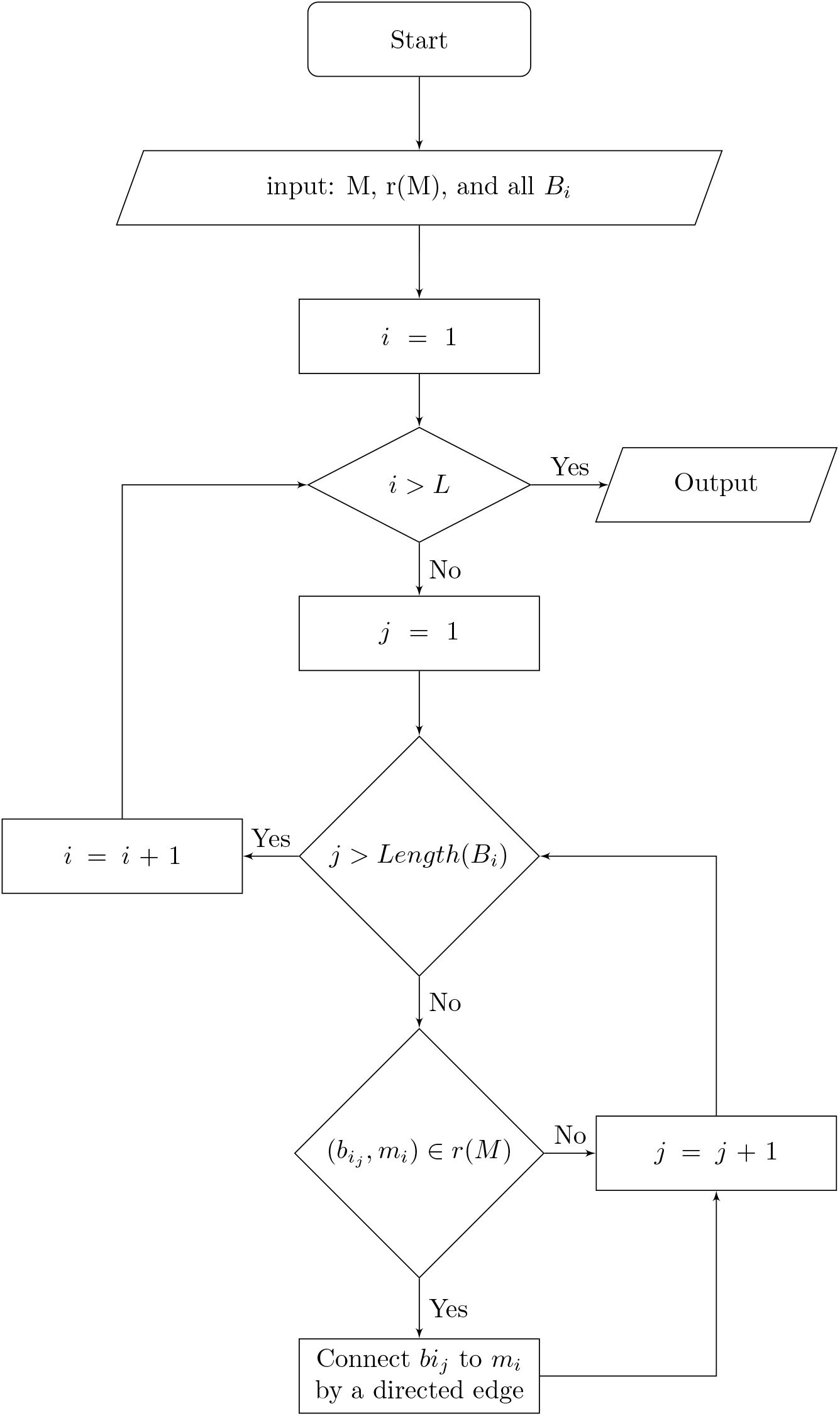

The output for both algorithms is given by identical pseudocode shown below.

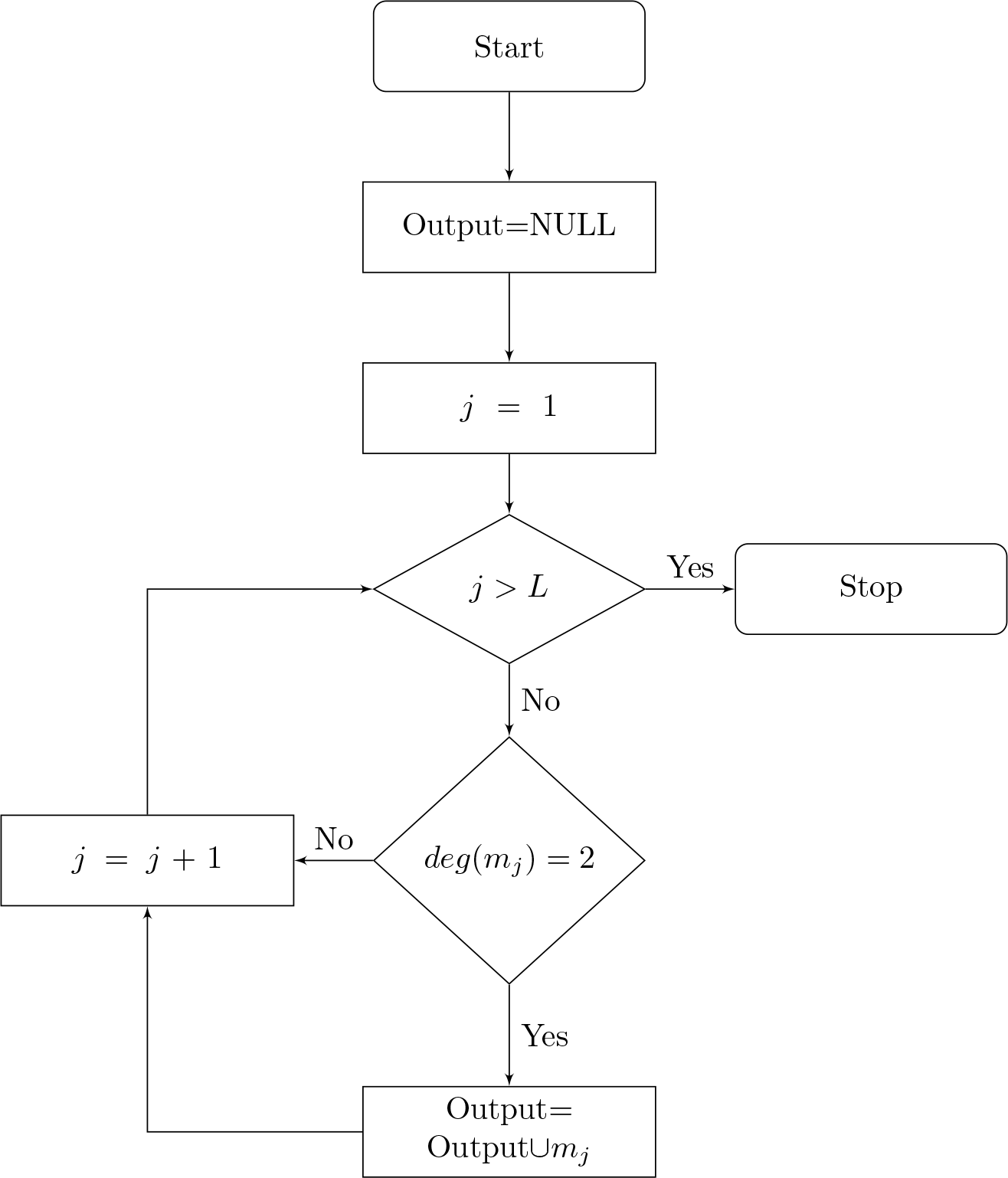

